# Accelerating genomic workflows using NVIDIA Parabricks

**DOI:** 10.1101/2022.07.20.498972

**Authors:** Kyle A. O’Connell, Zelaikha B. Yosufzai, Ross A. Campbell, Collin J. Lobb, Haley T. Engelken, Laura M. Gorrell, Thad B. Carlson, Josh J. Catana, Dina Mikdadi, Vivien R. Bonazzi, Juergen A. Klenk

**Affiliations:** Biomedical Data Science Team, Deloitte Consulting LLP, Arlington VA, 20057; Cloud Managed Services, Deloitte Consulting LLP, Detroit MI, 48226

**Keywords:** GPU acceleration, NVIDIA Parabricks, Cloud Computing, Amazon Web Services, Google Cloud Platform

## Abstract

**Background:** As genome sequencing becomes a more integral part of scientific research, government policy, and personalized medicine, the primary challenge for researchers is shifting from generating raw data to analyzing these vast datasets. Although much work has been done to reduce compute times using various configurations of traditional CPU computing infrastructures, Graphics Processing Units (GPUs) offer the opportunity to accelerate genomic workflows by several orders of magnitude. Here we benchmark one GPU-accelerated software suite called NVIDIA Parabricks on Amazon Web Services (AWS), Google Cloud Platform (GCP), and an NVIDIA DGX cluster. We benchmarked six variant calling pipelines, including two germline callers (HaplotypeCaller and DeepVariant) and four somatic callers (Mutect2, Muse, LoFreq, SomaticSniper).

**Results:** For germline callers, we achieved up to 65x acceleration, bringing HaplotypeCaller runtime down from 36 hours to 33 minutes on AWS, 35 minutes on GCP, and 24 minutes on the NVIDIA DGX. Somatic callers exhibited more variation between the number of GPUs and computing platforms. On cloud platforms, GPU-accelerated germline callers resulted in cost savings compared with CPU runs, whereas somatic callers were often more expensive than CPU runs because their GPU acceleration was not sufficient to overcome the increased GPU cost.

**Conclusions:** Germline variant callers scaled with the number of GPUs across platforms, whereas somatic variant callers exhibited more variation in the number of GPUs with the fastest runtimes, suggesting that these workflows are less GPU optimized and require benchmarking on the platform of choice before being deployed at production scales. Our study demonstrates that GPUs can be used to greatly accelerate genomic workflows, thus bringing closer to grasp urgent societal advances in the areas of biosurveillance and personalized medicine.

## BACKGROUND

As the cost of genome sequencing continues to decrease, genomic datasets grow in both size and generation speed (Langmead & Nellore, 2018). These processes will greatly enhance aims such as whole genome biosurveillance and personalized medicine (Nwadiugwu & Monteiro, 2022; Zhao et al., 2020). However, one challenge to attaining these goals is the computational burden of analyzing large amounts of genomic sequence data (Liu et al., 2014). Two trends (among others) are helping to ameliorate this burden. The first is the migration to the cloud for data analysis and storage, and the second is the use of Graphics Processing Units (GPUs) to accelerate data processing and analysis (Cole & Moore, 2018); (Franke & Crowgey, 2020). We address each of these trends in this article.

Cloud computing addresses many of the challenges associated with large whole genome sequencing projects, which can suffer from siloed data, long download times, and slow worlkflow runtimes (Tanjo et al., 2021). Several papers have reviewed the potential of cloud platforms for sequence data storage, sharing, and analysis (Augustyn et al., 2021; Cole & Moore, 2018; Grossman, 2019; Grzesik et al., 2021; Koppad et al., 2021; Langmead & Nellore, 2018; Leonard et al., 2019), thus here we focus on one cloud computing challenge, how to select the right compute configuration to optimize both cost and performance (Krissaane et al., 2020; Ray et al., 2021).

GPU acceleration in either a cloud or High Performance Computing (HPC) environment makes rapid genomic analysis possible at a scale previously not possible. While these are still early days for GPU-acceleration in the ‘omics fields, several studies have begun benchmarking various algorithmic and hardware configurations to find the balance between cost and performance. Franke & Crowgey, (2020) and Rosati, (2020) both benchmarked GATK HaplotypeCaller using the original CPU algorithm and the GPU-accelerated version from NVIDIA Clara™ Parabricks (https://www.parabricks.com/; hereafter Parabricks) on HPC platforms and found notable acceleration (8x and 21x speedups respectively) when using GPUs. They also inferred high concordance of SNP calls (∼99.5%) between the CPU and GPU algorithms suggesting no loss of accuracy with the GPU-configured algorithms, for both germline and somatic variant callers (*Benchmarking NVIDIA Clara Parabricks Somatic Variant Calling Pipeline on AWS*, 2022). Likewise, Zhang et al., (2021) introduced a new GPU-accelerated pipeline called BaseNumber, which achieved runtimes slightly faster than previous benchmarks using Parabricks.

While the aforementioned studies conducted benchmarking using on-premises computing clusters, some studies have begun benchmarking GPU-accelerated algorithms in the cloud. The Parabricks team at NVIDIA benchmarked GATK HaplotypeCaller using Parabricks on Amazon Web Services (AWS) and achieved runtimes as low as 28 minutes for a 30x genome with eight A100 NVIDIA GPUs (*Benchmarking the NVIDIA Clara Parabricks Germline Pipeline on AWS*, 2021), and speedups ranging from 4x to 42x for somatic callers (*Benchmarking NVIDIA Clara Parabricks Somatic Variant Calling Pipeline on AWS*, 2022). Relatedly, Krissaane et al., (2020) benchmarked GWAS workflows using Spark Clusters (not NVIDIA Parabricks) on both Google Cloud Platform (GCP) and Amazon Web Services (AWS) and found identical performance between cloud platforms. While these studies have shed light on the performance of GATK HaplotypeCaller using Parabricks, fewer studies have compared CPU and GPU performance for additional germline and somatic variant callers, or compared performance across AWS, GCP and an NVIDIA DGX cluster.

Here, we benchmark two germline variant callers and four somatic variant callers comparing traditional x86 CPU algorithms with GPU-accelerated algorithms implemented with NVIDIA Parabricks on AWS and GCP, and benchmark GPU-accelerated algorithms on an NVIDIA DGX cluster. In the case of GPU-accelerated algorithms, we compare 2, 4, and 8 GPU configurations. For germline callers, we observed speedups of up to 65x (GATK HaplotypeCaller) and found that performance scaled linearly with the number of GPUs. We also found that because GPUs run so quickly, researchers can save money by using them for germline variant callers. Alternatively, somatic variant callers achieved speedups up to 56.8x for the Mutect2 algorithm, but surprisingly, did not scale linearly with the number of GPUs, emphasizing the need for algorithmic benchmarking before embarking on large-scale projects.

## RESULTS

### CPU baseline across cloud platforms

CPU machine performance varied considerably between AWS/GCP for most analyses. For germline analyses, GCP performed faster for DeepVariant (18.8 hrs) compared with AWS (22 hrs), whereas AWS performed faster for HaplotypeCaller (36.2 hrs) compared with GCP (38.8 hrs; Table 1, Fig. 1). Somatic runtimes favored AWS, with the exception of Mutect2, where GCP ran in 8.1 hrs compared with 16.9 hrs on AWS (Table 1, Fig. 1).

**Table 1:**
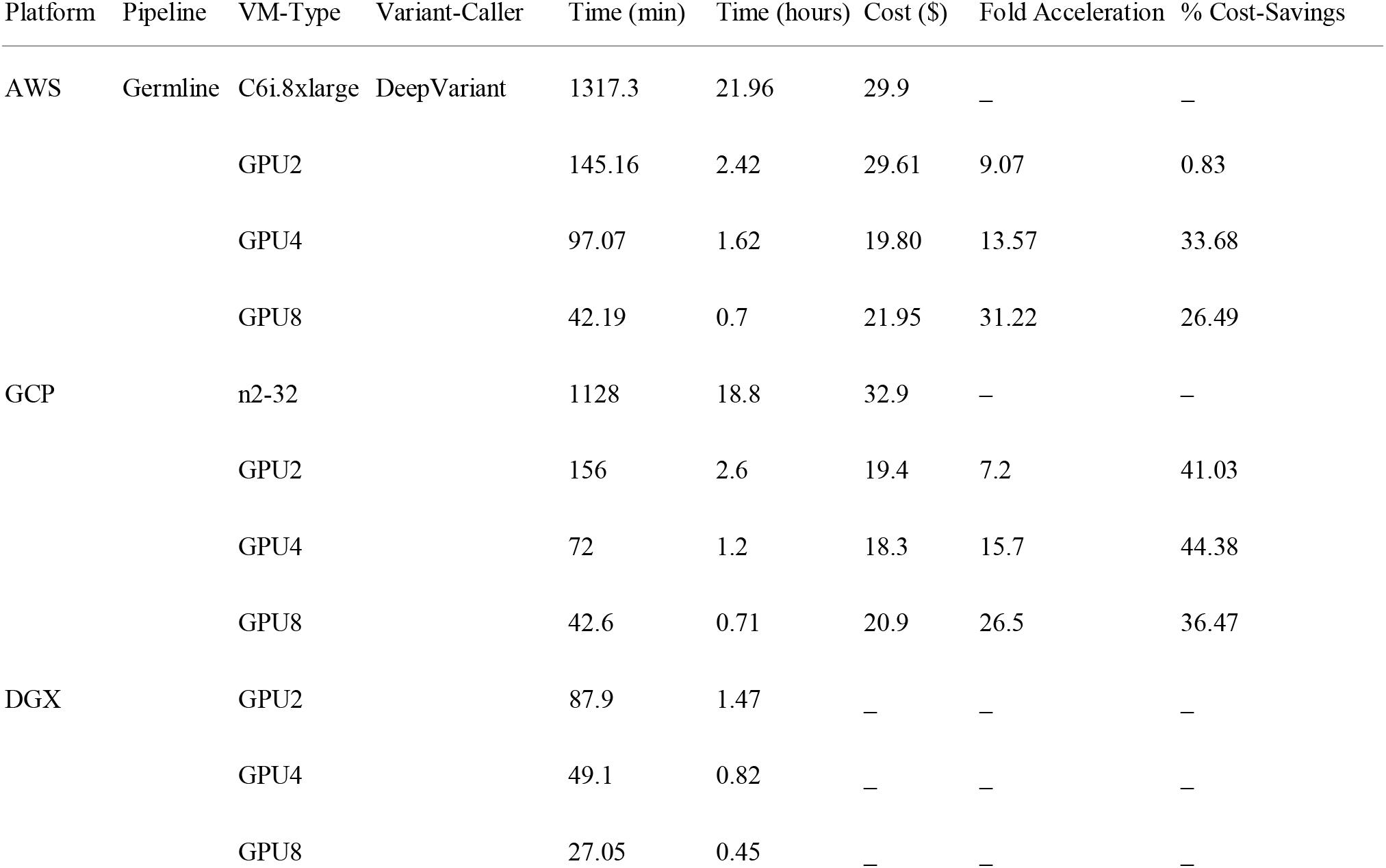
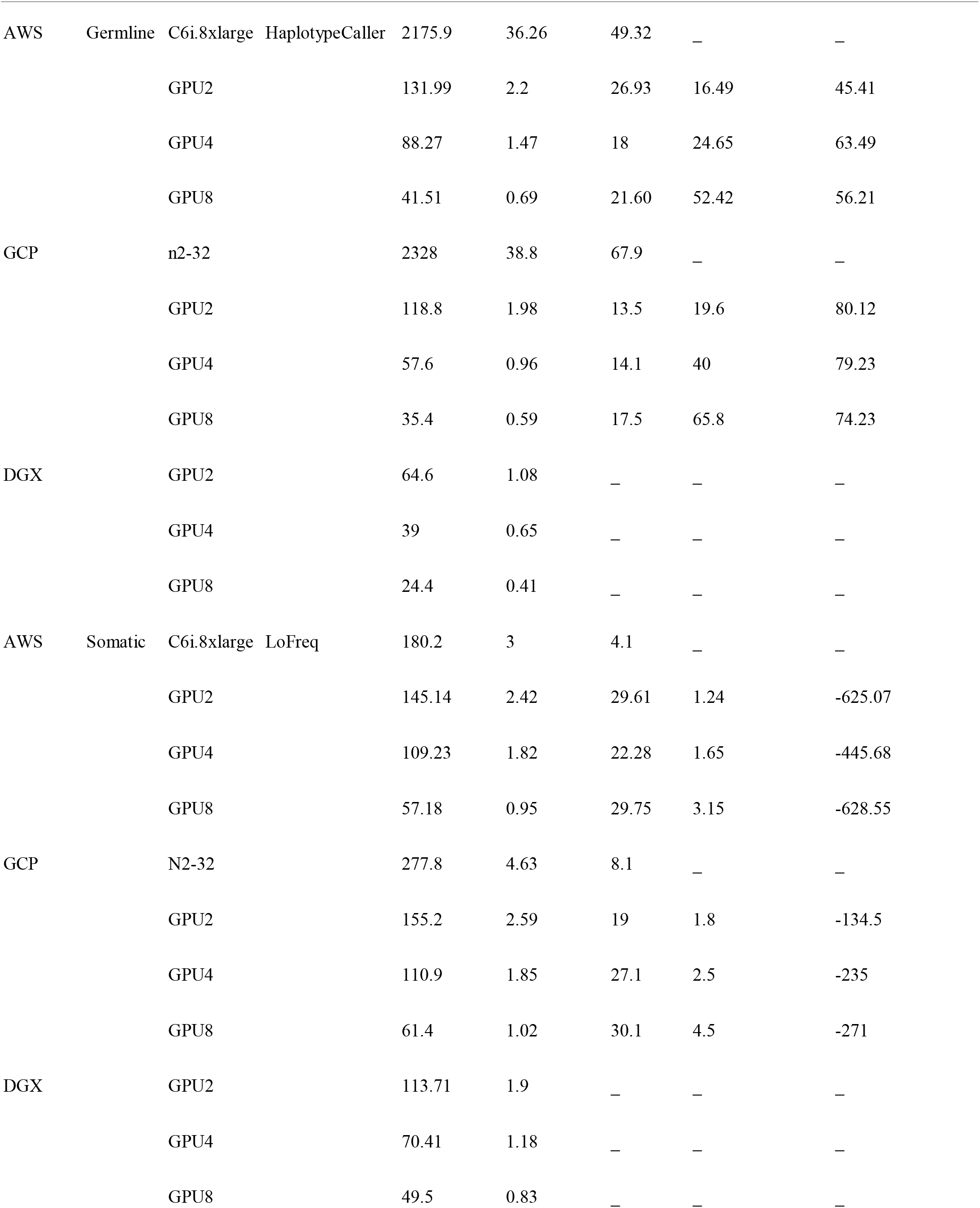
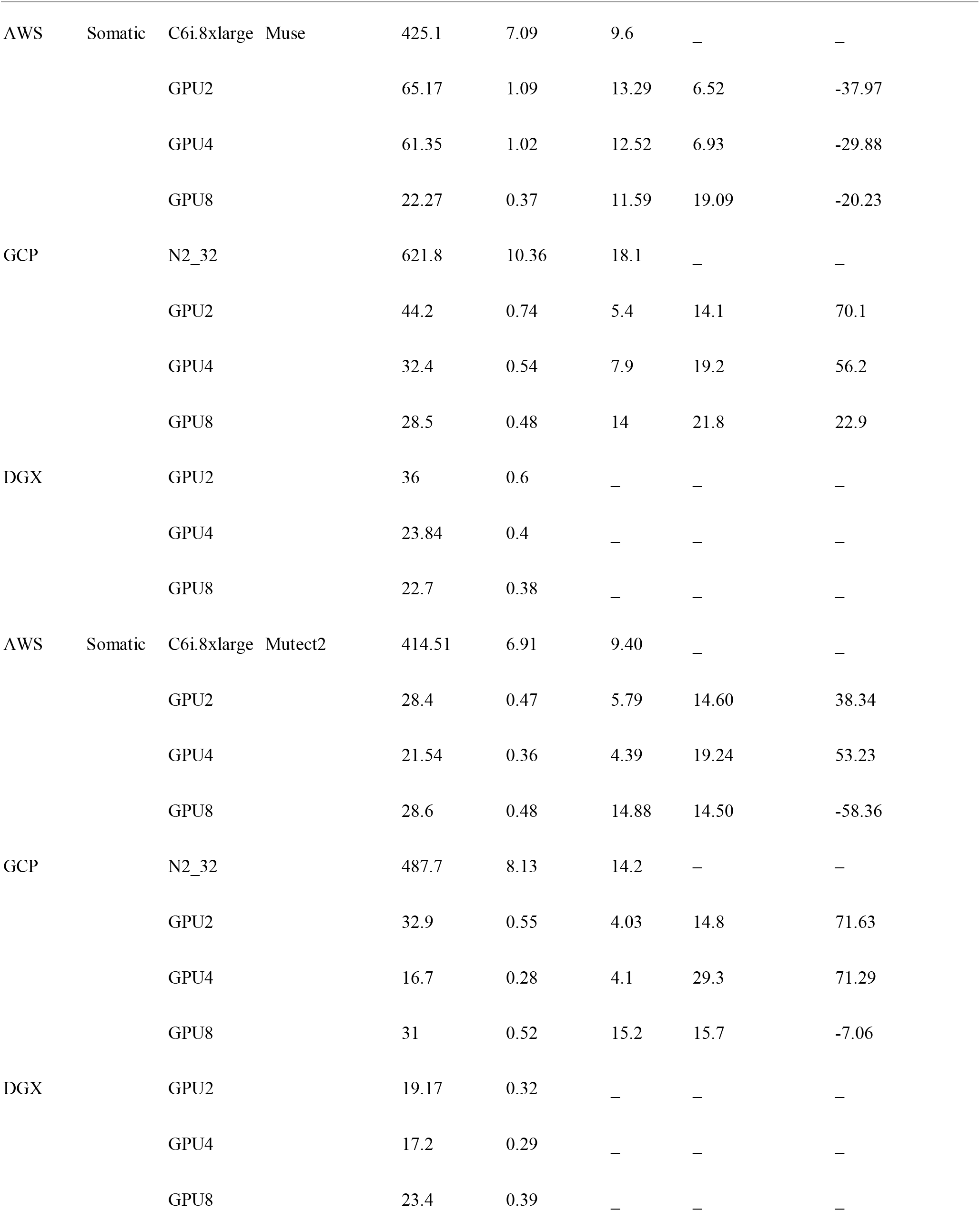
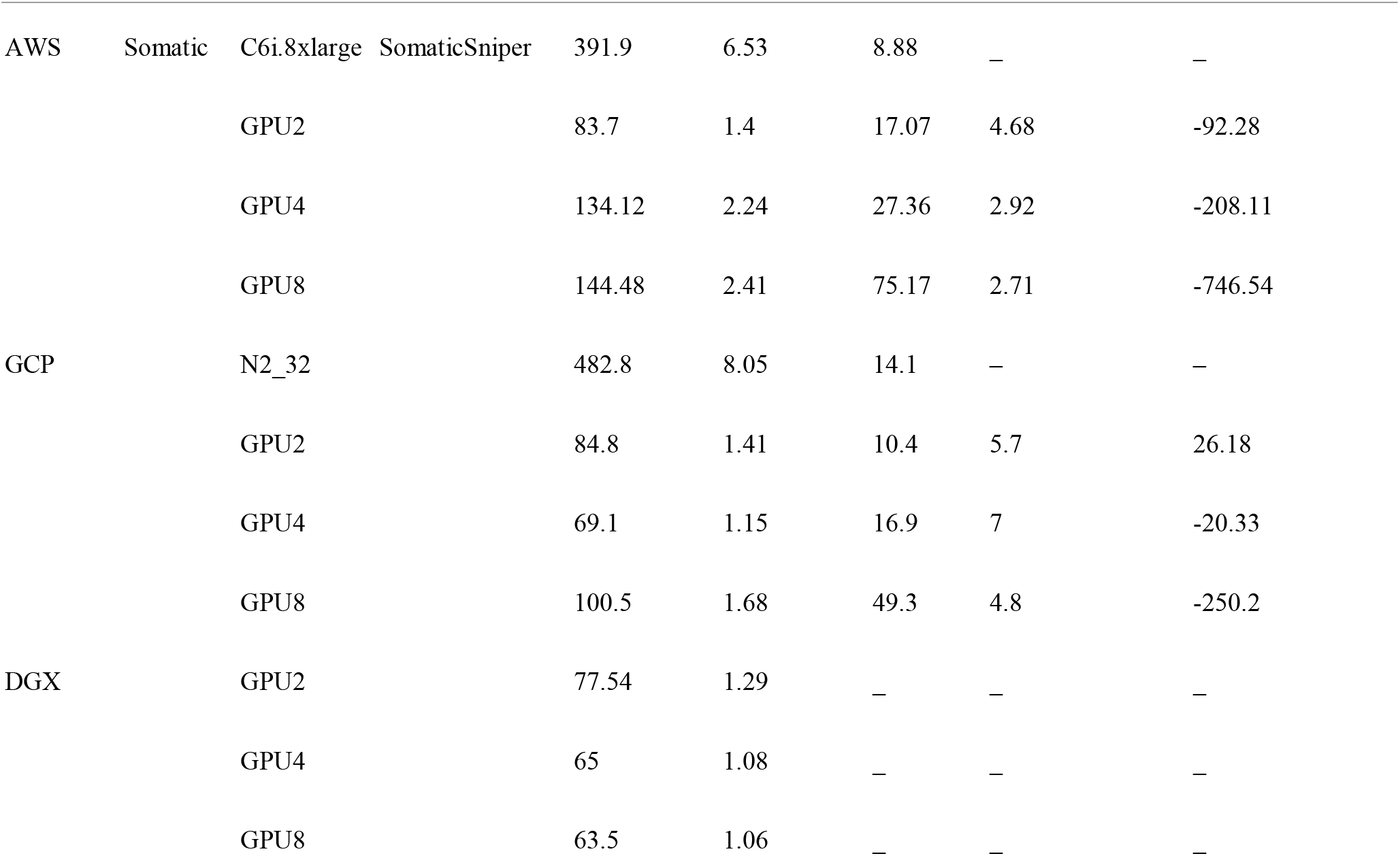
Results of benchmarking for AWS, GCP and NVIDIA DGX workflow runs. AWS results presented here are for the p3 family with the NVIDIA Tesla V100 GPU, results for the p4 family with the A100 GPU are shown in Table S1

**Figure 1.**
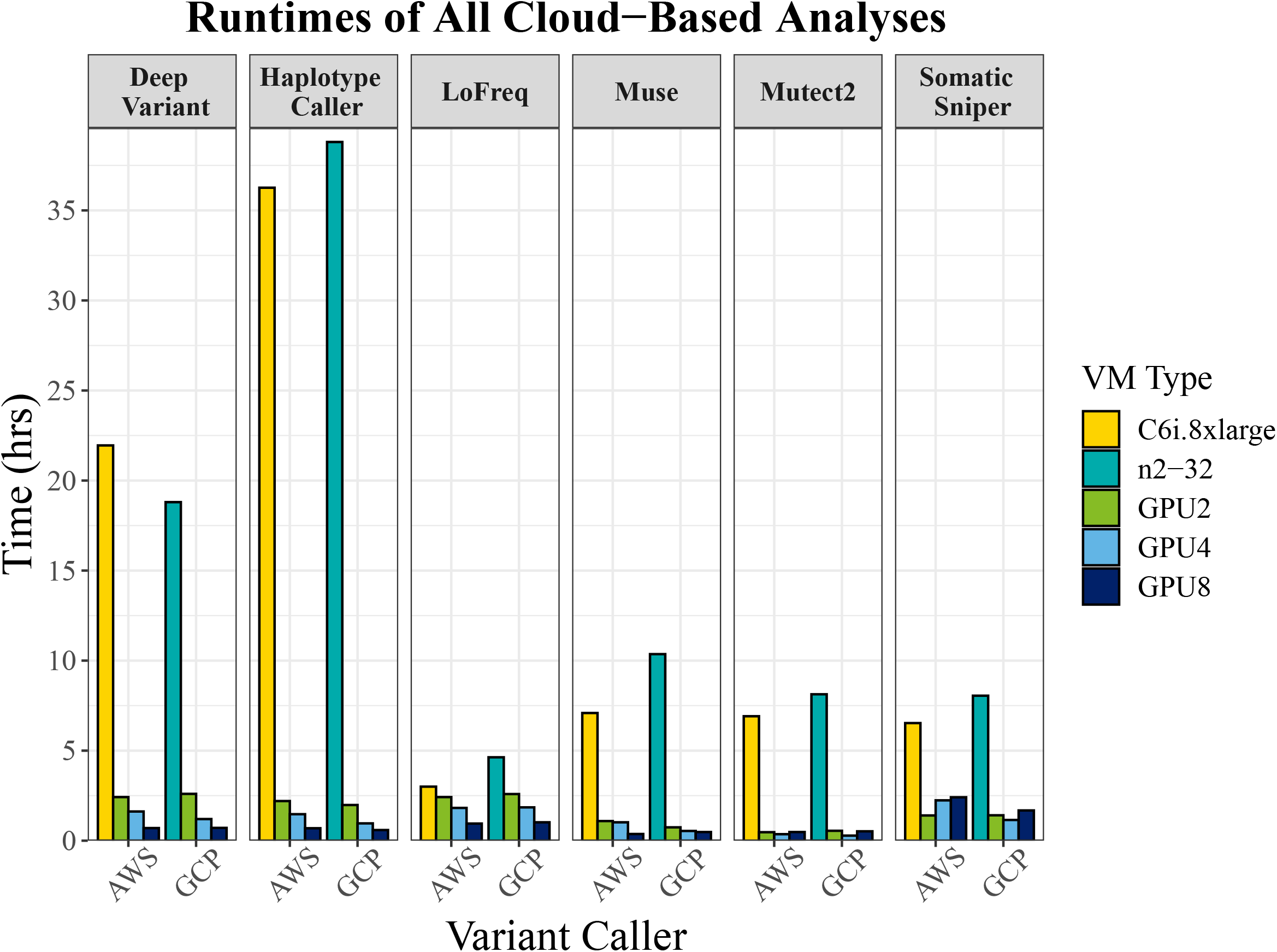
Comparison of execution times of variant calling algorithms on CPU and GPU environments between AWS and GCP. A 32 vCPU machine with the latest processors was used for CPU benchmarking on both cloud platforms. Here we show results for varying numbers of NVIDIA Tesla V100 GPUs running the Parabricks bioinformatics suite for AWS, and NVIDIA Tesla A100 GPUs for GCP.

### GPU performance across cloud platforms

For germline callers, 8-GPU runtimes were below 45 minutes for HaplotypeCaller and DeepVariant across both cloud platforms. On AWS, we observed faster runtimes for the A100 compared with the V100 GPU machines (p4 vs p3 machine families), but the differences with 8 GPUs, where the number of CPUs were equal, were small for most workflows. Further, comparisons between the 2 and 4 A100 GPU machines on GCP/AWS was not precise because we were unable to limit the number of CPUs available for all workflows, thus differences in times between the two cloud platforms were biased towards AWS for some algorithms. Although the two germline workflows scaled linearly with the number of GPUs (Fig. 2), somatic callers ran faster with 4 vs. 8 GPUs for

**Figure 2.**
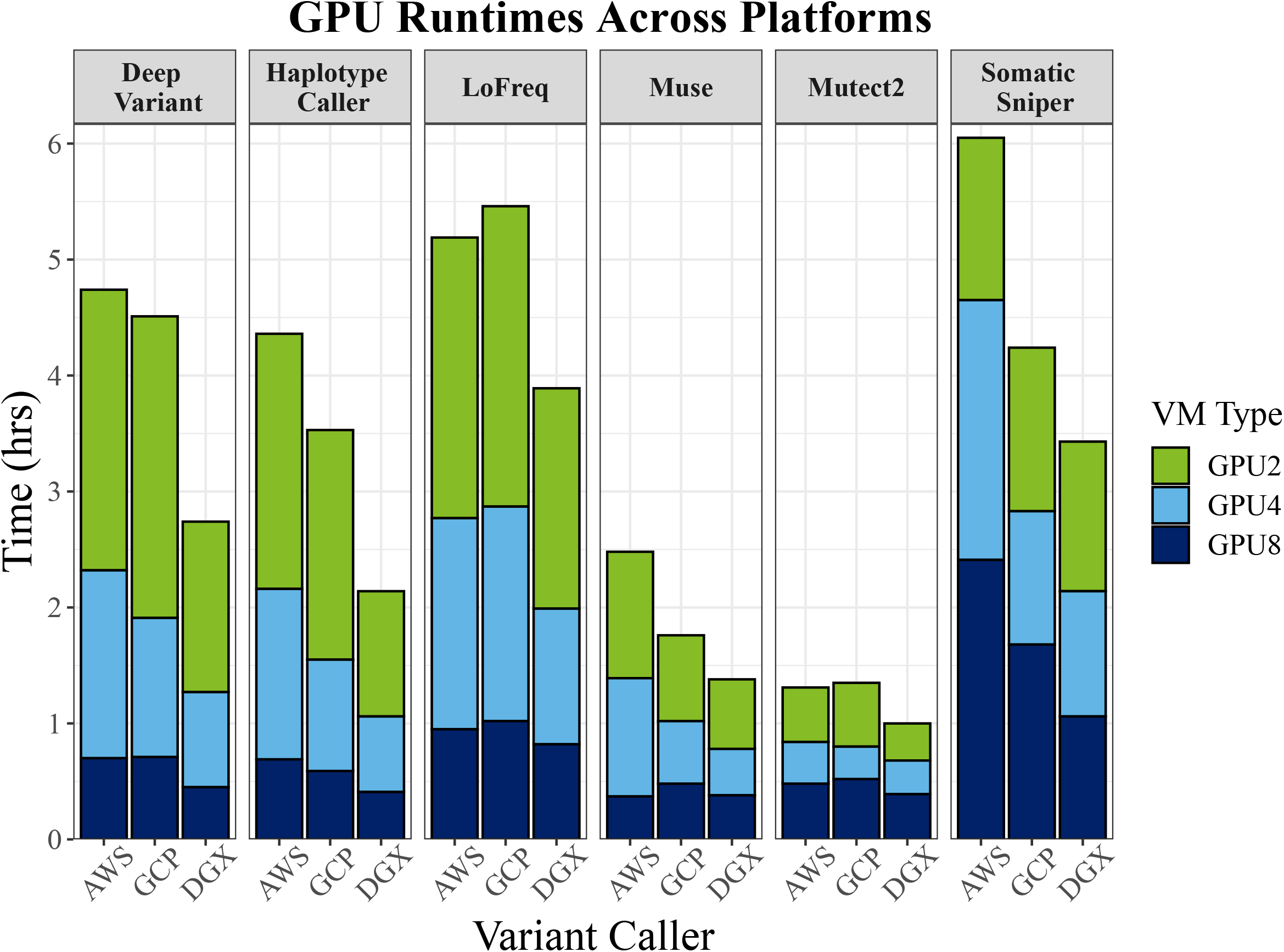
GPU benchmarking results for NVIDIA Tesla GPUs. On GCP and the DGX results are shown for A100 GPUs, whereas AWS results are shown for the V100 GPU runs.

Muse on AWS (but not GCP), Mutect2 on AWS and GCP, and for SomaticSniper on AWS and GCP (Fig. 2; S1). Compared with the CPU baselines, GPU runs on AWS (with A100 GPU) led to acceleration of HaplotypeCaller up to 65.1x, DeepVariant up to 30.7x, Mutect2 up to 56.8x, SomaticSniper up to 7.7x, Muse up to 18.9x, and Lofreq up to 3.7x (Table 1). On GCP, GPUs resulted in acceleration of HaplotypeCaller up to 65.8x, DeepVariant up to 26.5x, Mutect2 up to 29.3x, SomaticSniper up to 7.0x, Muse up to 21.8x, and LoFreq up to 4.5x.

Although GPU machines are much more expensive than CPU machines, the accelerated runtimes result in cost savings for most algorithms (Fig. 4). Leveraging GPUs on AWS with the A100 machine resulted in cost savings up to 63% for HaplotypeCaller with 8 GPUs, 33% for DeepVariant with 4 GPUs, and up to 57.6% for Mutect2 with 4 GPUs. Using the A100 GPU machine resulted in even greater savings of 63% for HaplotypeCaller with 4 GPUs, 21% for DeepVariant with 8 GPUs, and 80% for Mutect2 with 4 GPUs (Table S1).

On GCP GPU runs resulted in cost savings of up to 80.1% for HaplotypeCaller with 2 GPUs, 44.4% for DeepVariant with 4 GPUs, 71.6% for Mutect2 with 4 GPUs, 26.2% for SomaticSniper with 2 GPUs, and up to 70.1% for Muse with 2 GPUs. However, on both platforms, algorithms that were not well optimized actually cost much more to run with GPUs rather than CPUs because the difference in runtimes was not enough to offset the extra GPU cost (Fig. 4; S4). For example, CPU runs of LoFreq cost less than $10/sample to run on both platforms, but as much as $30 with GPUs (Fig. S2). Likewise, CPU runs of Somatic Sniper cost less than $15 per sample on both platforms, but as much as $75 on AWS with 8 GPUs.

**Figure 3.**
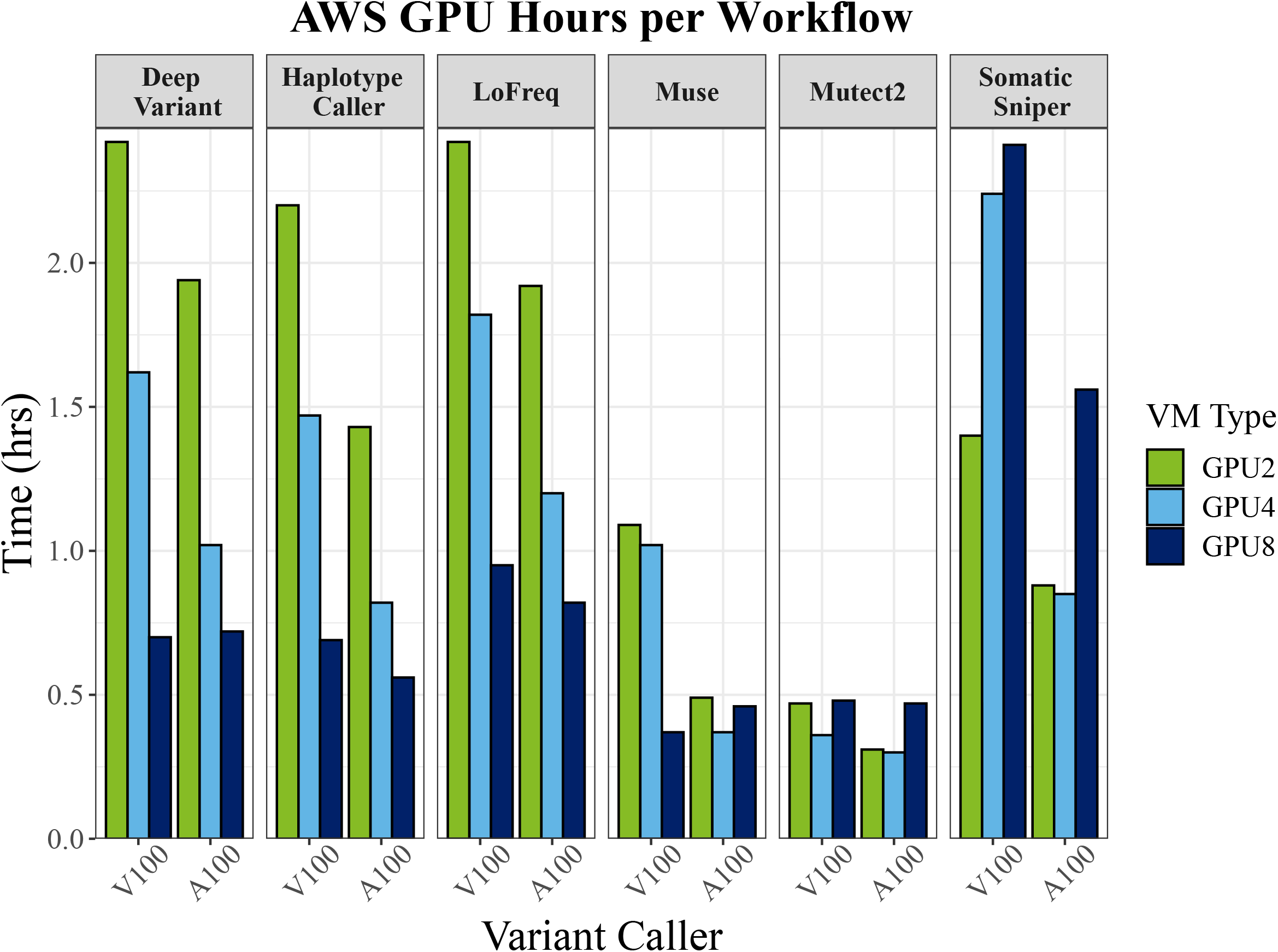
Comparison of runtimes between V100 and A100 GPU machines on AWS

**Figure 4.**
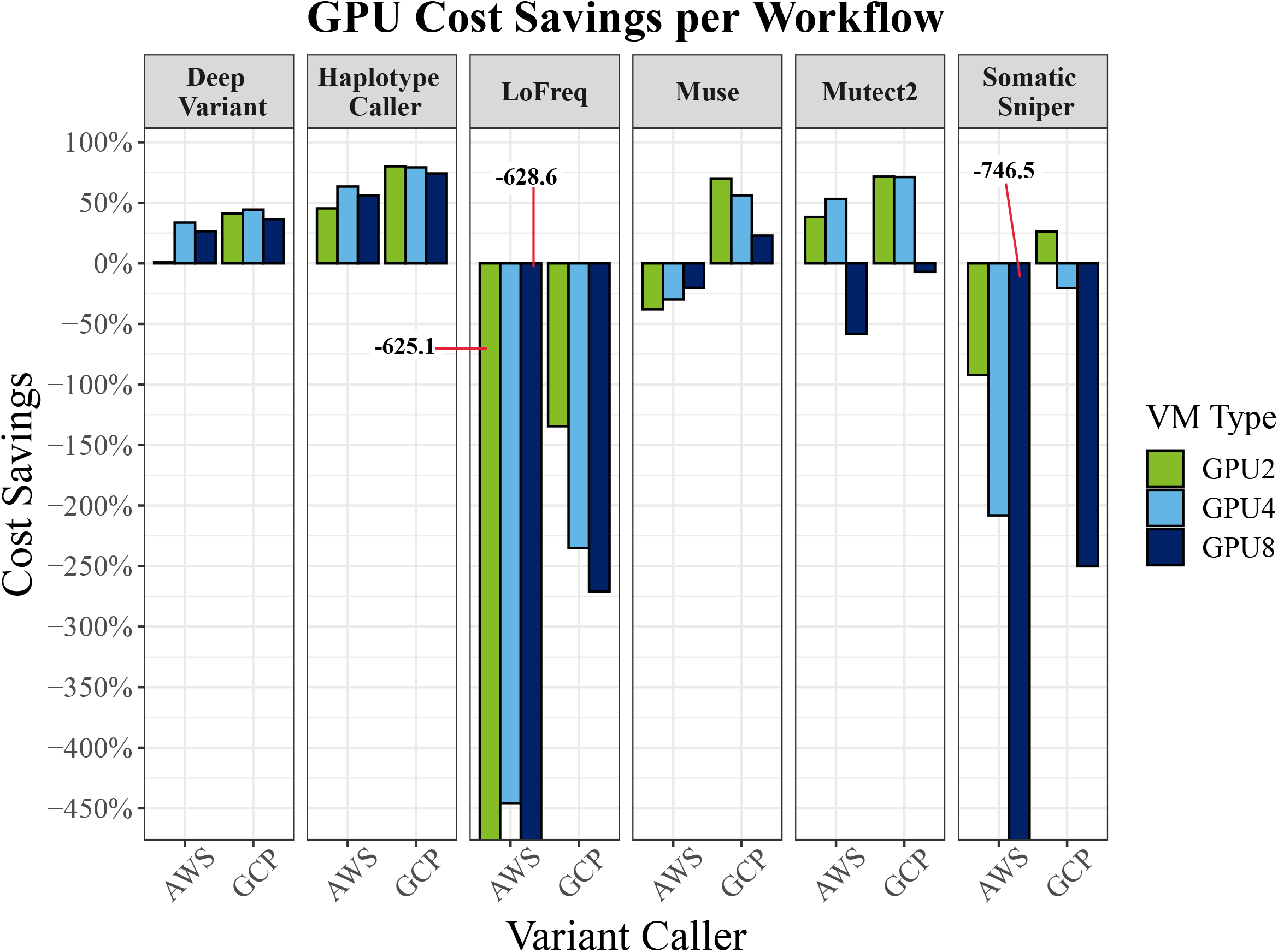
Comparison of AWS (V100 GPU machine) vs. GCP GPU cost savings per variant caller. Percentage of total cost savings shows a majority of higher cost savings using GPUs in algorithms optimized for GPU-acceleration, but losses when algorithms are not well optimized

For well optimized algorithms, results varied between variant callers on which numbers of GPUs were the fastest (ranging from 2–8); subsequently cost savings reflect a balance between speed and cost of a particular machine type that is not consistent between algorithms or cloud providers. For example, A100 GPU runs were expensive on AWS because the p4d.24xlarge machine type on demand price is $32.8/hr, whereas the A100 machine type ranges from $12.24/hr for a 4 GPU machine, to $24.5/hr for an 8 GPU machine. On GCP, the a2-highgpu machine types range from $7.4/hr (2 GPUs) to $29.4.00/hr (8 GPUs). Alternatively, CPU runs were slightly cheaper on AWS with an on demand price of $1.36/hr compared with $1.75 on GCP. Prices here are given for the northern Virginia region calculated (at the time of writing) using the pricing calculators from the respective cloud service providers. As time goes on, these machine types will likely become less expensive with greater adoption.

### GPU performance on the DGX

Germline workflows ran considerably faster on the DGX than on the cloud platforms, with HaplotypeCaller finishing in 24.4 min and DeepVariant finishing in 27.1 min with 8 GPUs (Fig. 2; S1). Somatic variant callers were not faster in most cases than the cloud platforms, and in one case, ran slower than on the cloud (Somatic Sniper; Fig. 2; S1). Interestingly, the pattern we observed in the cloud where the 4 GPU runtimes were the fastest for Muse and Somatic Sniper did not manifest on the DGX, where the 8 GPU runs were the fastest for all algorithms, with the exception of Mutect2 (Fig. 2; S1). For Mutect2, the 4 GPU run was still the fastest on the DGX, but the 8 GPU run was faster on the DGX than on both AWS/GCP (Fig. S1).

We also tested the effect of CPU number on performance of GPU runs. On AWS and GCP the GPU machine types are preconfigured with 12 CPUs/1 GPU, but on the DGX we were able to modify the number of CPUs for each run. We found that adding CPUs does decrease runtimes (increase performance), but that reduction of runtimes plateaued after 48 CPUs (Fig. S5).

## DISCUSSION

The acceleration provided by GPU-accelerated algorithms confers several advantages to researchers. First, GPU-acceleration enables researchers to rapidly run multiple algorithms (Crowgey et al., 2021). Different variant callers exhibit biases leading to slightly different variant calls (Zhao et al., 2020). Combining calls across algorithms can lead to higher accuracy, albeit with a slightly higher type one error. Future studies could compare false positive and negative rates for different strategies of combining calls across algorithms such as majority rule vs. consensus site calls. Another advantage of GPU-accelerated genomic workflows is that they allow researchers to process more samples with a fixed budget. Academic research programs are often constrained by limited funding; the use GPU-acceleration may allow researchers to reduce compute costs (and labor overhead) and thus process more samples for the same amount of money. Finally, GPU-accelerated algorithms enable near-real-time decision making. Pathogen biosurveillance benefits from rapid data processing to identify novel pathogens and allow policymakers to act before an outbreak spreads (Gardy & Loman, 2018).

### Cloud platform considerations

#### CPU only runs

As more research programs migrate to cloud platforms, researchers will need to make decisions about which platform provides the most advantages for both performance and cost considerations. CPU runs were faster on the AWS c6i.8xlarge machine than on the GCP n2-32 for four algorithms, while DeepVariant and Mutect2 ran faster on GCP (Fig. 1). Both of these machine types use the newest Intel Xeon Scalable processors (Ice Lake), but seem to have inherent differences that would be difficult to identify without benchmarking particular algorithms as we have done here. Regardless of cloud platform however, past work within our research group showed that reduced runtimes driven by using the latest CPU processors outweighs the increased per second cost (TC unpublished).

Another consideration that researchers should be aware of in the near term is that AWS is migrating to newer ARM-based machine types, rather than x86 architectures. We had trouble installing existing software on the ARM-based machines, and thus used the c6i.8xlarge machine. This could present challenges for researchers in the future on AWS as the platform migrates more machine types to ARM-based architectures, necessitating the rewriting and/or compiling of common software. On GCP, we chose the N2 machine family as a balance between performance and cost. GCP does offer the compute-optimized C2 machine family, which may run faster than the N2 machines, but we did not benchmark those machines here.

#### GPU considerations on the cloud

For germline workflows, AWS and GCP performed very similarly for both speed and cost when using 8 A100 GPUs, although the 2 and 4 GPUs runs exhibited more variation (Fig. 2,4). In an effort to quantify the balance between cost and performance on each cloud platform, we calculated a cost ratio metric by dividing the cost of the workflow by the xSpeedup for a GPU run when compared to the CPU run for that workflow. Thus, a lower cost ratio indicates a better value for a given GPU configuration (Table 1; Fig. 5). For the germline variant callers, the best cost ratio on both platforms used 8 GPUs, and the ratio for AWS and GCP was similar enough that we feel it should not impact the choice between cloud providers. For somatic variant workflows, the best cost ratio was usually 2–4 GPUs, as these workflows were not well optimized to use 8 GPUs on the cloud. Further, because LoFreq and Somatic Sniper were not very accelerated with Parabricks, their high cost ratio suggests that it is not worth the extra cost to run these workflows using GPUs. It should be noted that we only benchmarked using on demand instances, and bioinformaticians could save additional costs by leveraging spot instances.

**Figure 5.**
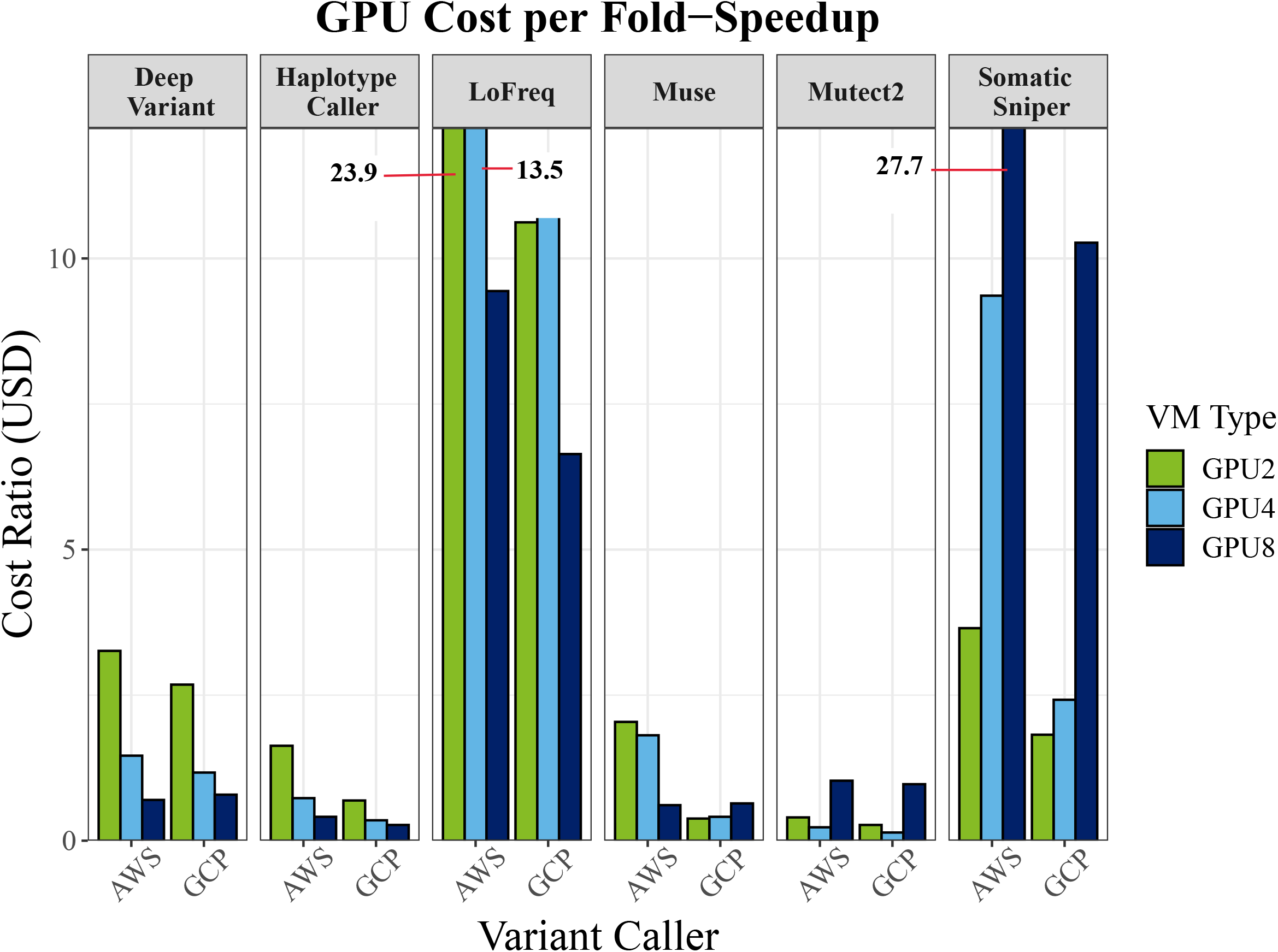
Comparison of AWS V100 vs. GCP A100 GPU cost ratio per variant caller. Cost ratio being the ratio between cost per hour and fold speed-up. Cost per fold-speedup shows the benefit of harnessing GPU over CPU in select algorithms, while other algorithms are more cost-efficient with CPUs.

GPU-accelerated bioinformatic workflows are still relatively new to the cloud, and as such, not all tools are readily available everywhere. For example, at the time of writing, Parabricks did not offer a Marketplace solution for GCP, although their team was working on releasing one (G. Burnett *pers. comm*). Likewise, the Marketplace solution on AWS offered a user-friendly way to access the Parabricks software suite without purchasing an annual license, but this machine image did not support the p4 machine family with the A100 GPUs. Nonetheless, although we were able to install Parabricks on the A100 machine on AWS, this machine type was not readily available (at the time of writing) in most regions, and it was difficult to procure this machine type to conduct our benchmarking. Perhaps using spot instances would have been a better solution for these difficult to procure machine types. Finally, we observed some decreases in runtime between the A100 and V100 GPU machines on AWS (Fig. 3). However, differences were relatively minor when using 8 GPUs – less than a minute for DeepVariant and eight minutes for HaplotypeCaller. As long as the A100 machine type is difficult to obtain and is not available with the Marketplace machine image, we recommend using the V100 GPU machine without much cost to performance (Table 1, S1; Fig. S3).

### On-premises computing clusters

For a myriad of reasons, some bioinformatic analyses will not migrate to the cloud, thus requiring on-premises infrastructure. Although not every institution will have a DGX cluster with A100 GPUs available, we show here that Parabricks runs well in an on-premises environment. For those looking to achieve the fastest possible runtimes in a production environment, the DGX ran considerably faster than AWS or GCP for germline callers, reducing runtimes for HaplotypeCaller by 8 min and DeepVariant by 15 min, differences that could be significant at large enough scales. We attribute these differences to the network communication between GPUs and CPUs on the machines, which is better optimized on the DGX compared with cloud-based instances, where GPUs may not be located in as close of proximity

## CONCLUSIONS

We found that germline variant callers were well optimized with Parabricks and that GPU-accelerated workflows can result in substantial savings of both time and costs. Alternatively, somatic callers were accelerated, but exhibited substantial variation between algorithms, number of GPUs, and computing platform, suggesting that benchmarking algorithms with a reduced dataset is important before scaling up to an entire study. Though early days for GPU-accelerated bioinformatic pipelines, ever faster computing processors bring us closer to important societal aims such as tracking pathogens in near real-time to monitor emerging pandemics or enabling milestones in the field of personalized medicine.

## MATERIALS AND METHODS

### Sampling and Algorithms

We benchmarked six variant callers for CPU and GPU speed and cost. We conducted all benchmarking on the individual ‘HG002’ from the Genome in a Bottle Consortium (Krusche et al., 2019; Zook et al., 2016) hosted by the National Institute of Standards and Technology, and made available as part of the Precision FDA Truth Challenge V2 (https://precision.fda.gov/challenges/10). We downsampled the fastq files to 30x coverage using Samtools (Li et al., 2009). We used Grch38 as our reference genome downloaded from the GATK Reference Bundle. Our germline variant calling pipeline evaluated two germline variant callers: HaplotypeCaller (Poplin, Chang, et al., 2018; Van der Auwera & O’Connor, 2020) and DeepVariant (Poplin, Ruano-Rubio, et al., 2018). GPU benchmarking used Parabricks. For germline callers we used ‘Germline Pipeline’ for GATK HaplotypeCaller, and for DeepVariant we used ‘DeepVariant Germline Pipeline‘. Each of these pipelines take fastq files as inputs and output unfiltered variant call format (VCF) files. CPU benchmarking was conducted by writing custom workflows using Snakemake (Mölder et al., 2021), following best practices for each tool and exactly matching the workflows used by Parabricks (Data Accessibility).

Our somatic variant calling pipeline evaluated four somatic variant callers: Mutect2 (Van der Auwera & O’Connor, 2020), SomaticSniper (Larson et al., 2012), Muse (Fan et al., 2016), and LoFreq (Wilm et al., 2012). We generated synthetic somatic tumor data using SomatoSim (Hawari et al., 2021). We added 198 single nucleotide polymorphisms (SNPs) at random variant allele frequencies ranging from 0.001 to 0.4 (randomly generated using custom python scripts). Sites were selected from the ICGC Data Portal ovarian cancer patient DO32536 (https://dcc.icgc.org/donors/DO32536?mutations=%7B%22size%22:50,%22from%22:151%7D). We used the BAM file from the HaplotypeCaller pipeline (i.e., MarkDuplicates, BaseRecalibration, and ApplyBQSR were run prior to the mutation process) as the input for SomatoSim. For somatic variant callers, we used the Parabricks variant caller scripts (‘mutectcaller’, ‘somaticsniper_workflow’, ‘muse’, ‘lofreq’) which take BAM files as input and output VCF files. Each Parabricks tool was compared to a compatible CPU command as listed in the Parabricks 3.7 documentation. We used Snakemake scripts as described for germline callers. For benchmarking of MuSE, we used version v2.0 and set the number of threads to 1 to replicate MuSE v1.0 lack of parallel computing because of version conflicts with MuSE v1 in our compute environment. We created a conda environment before running each workflow because we found that using the ‘--with conda’ flag in Snakemake dramatically increased run times. Complete workflows are described in the Supporting Information and all scripts necessary to repeat our analyses are available at (https://github.com/kyleoconnell/gpu-acclerated-genomics).

### GCP Configuration

Benchmarking on GCP leveraged virtual machines that were launched programmatically for CPU machines, or manually for GPU machines. CPU workflows used the ‘n2-standard-32’ machine type with Intel Xeon Cascade Lake processors with 32 vCPUs and128 GB of RAM. We assigned 1 TB of EBS storage to our instance. We launched these machines using a startup script that installed the conda environment, then ran the snakemake workflows. All data was already loaded on a machine image, and runtimes were concatenated from each snakemake rule using a custom script. We also benchmarked the older generation E2 family of processors, but found the run times to be much slower and thus only present the results for N2 processors here. GPU benchmarking on GCP used the accelerator optimized a2-highgpu machine types with two A100 GPUs, 24 vCPUs and 170GB RAM, four A100 GPUs with 48 vCPUs and 340 GB RAM, and eight A100 GPUs with 96 vCPUs and 680 GB RAM. One virtual machine was utilized with 4 TB storage, which we stopped and resized between runs.

### AWS Configuration

Benchmarking on AWS also used multiple virtual machines for CPU and GPU benchmarking. CPU benchmarking used the C6i.8xlarge machine type, which has a 3rd generation Intel Xeon Scalable processor with 32 vCPUs and 64 GiB RAM. We assigned 800 GB of EBS storage to our instance. We did some preliminary testing with the new ARM-based processors but had issues with installing several of the dependencies (particularly with mamba/conda), suggesting that a migration to ARM-based processors may prove problematic for bioinformatics in the cloud.

We benchmarked two GPU machine families. First, we benchmarked the p4 machine family which is similar to GCP a2-highgpu machines utilizing the latest NVIDIA A100 Tensor Core GPUs with 8 GPUs with 96 vCPUs and 1152 GiB RAM. AWS currently only has one machine type with A100 GPUs, the p4d.24xlarge, which only runs with 8 GPUs. To ensure consistency with GCP, we ran the 8 GPU machine, but specified the number of GPUs to use in our Parabricks commands for the smaller numbers of GPU runs. Because this machine type was not compatible with the marketplace image (see below) we installed Parabricks manually using scripts provided by NVIDIA. When possible (--cpu flag available) we limited the number of CPUs available with the p4 machine, but some analysis used more CPUs on AWS than on GCP.

To compare GPU and CPU configurations directly with GCP, we further benchmarked the p3 machine family using the ‘NVIDIA Clara Parabricks Pipelines’ AWS Marketplace image. At the time of writing the image supported V100 GPUs (but not A100 GPUs), which are an older model of Tensor Core GPU, on machine types p3.8xlarge with 4 GPUs and p3dn.24xlarge with 8 GPUs. The Marketplace image also had Parabricks preinstalled at a cost of $0.30 for the software. This configuration allowed us to directly compare 4, and 8 GPU machines with equal CPU numbers between AWS and GCP. Again, we limited the number of CPUs available to the 2 GPU runs when possible. After we finished our analyses, NVIDIA wrote a helpful somatic benchmarking guide (https://github.com/clara-parabricks/NVIDIA-Clara-Parabricks-Somatic-Variant-Calling-AWS-Blog).

### DGX Configuration

We also conducted GPU benchmarking on an NVIDIA DGX Cluster (DGX SuperPOD), which is a computing cluster with six DGX A100s, each of which contains eight NVIDIA A100 GPUs. Although the cluster technically has 48 A100 GPUs available, Parabricks is only able to run on a single DGX A100 system, thus limiting any Parabricks analyses to 8 GPUs. Jobs were launched using a Kubernetes-based scheduler, allocating a max memory of 300 GB, and matching the GPU and CPU configurations of the GCP/AWS runs, with the exception of GATK HaplotypeCaller. For this workflow, we benchmarked times for 8 GPUs using 24, 48, 96, and 124 CPUs.

## DECLARATIONS

### Availability of Data and Materials

The dataset(s) supporting the conclusions of this article is(are) available in the GitHub at https://github.com/kyleoconnell/gpu-acclerated-genomics.

### Author’s Contributions

KAO, CJL, TBC, DM, VRB, and JAK conceived the study. KAO, ZBY, RAC, and CJL designed the study. KAO, ZBY, RAC, and CJL ran cloud-based analyses. KAO and JJC ran DGX analyses. KAO, ZBY and HTE wrote the manuscript, and all authors read and approved of the text.

### Competing Interests

Deloitte Consulting LLP. and NVIDIA are alliance partners.

## Notes

### Competing Interest Statement

The authors have declared no competing interest.

## REFERENCES

Augustyn, D. R., Wyciślik, Ł., & Mrozek, D. (2021). Perspectives of using Cloud computing in integrative analysis of multi-omics data. Briefings in Functional Genomics, 20(4), 198– 206. https://doi.org/10.1093/bfgp/elab007

Benchmarking NVIDIA Clara Parabricks Somatic Variant Calling Pipeline on AWS. (2022, April 20). Amazon Web Services. https://aws.amazon.com/blogs/hpc/benchmarking-nvidia-clara-parabricks-somatic-variant-calling-pipeline-on-aws/

Benchmarking NVIDIA Clara Parabricks Somatic Variant Calling Pipeline on AWS. (2022, May 10). HPCwire. https://www.hpcwire.com/solution_content/aws/benchmarking-nvidia-clara-parabricks-somatic-variant-calling-pipeline-on-aws/

Benchmarking the NVIDIA Clara Parabricks germline pipeline on AWS. (2021, November 23). Amazon Web Services. https://aws.amazon.com/blogs/hpc/benchmarking-the-nvidia-clara-parabricks-germline-pipeline-on-aws/

Cole, B. S., & Moore, J. H. (2018). Eleven quick tips for architecting biomedical informatics workflows with cloud computing. PLOS Computational Biology, 14(3), e1005994. https://doi.org/10.1371/journal.pcbi.1005994

Crowgey, E. L., Vats, P., Franke, K., Burnett, G., Sethia, A., Harkins, T., & Druley, T. E. (2021). Enhanced processing of genomic sequencing data for pediatric cancers: GPUs and machine learning techniques for variant detection. Cancer Research, 81(13_Supplement), 165. https://doi.org/10.1158/1538-7445.AM2021-165

Franke, K. R., & Crowgey, E. L. (2020). Accelerating next generation sequencing data analysis: An evaluation of optimized best practices for Genome Analysis Toolkit algorithms. Genomics & Informatics, 18(1), e10. https://doi.org/10.5808/GI.2020.18.1.e10

Gardy, J. L., & Loman, N. J. (2018). Towards a genomics-informed, real-time, global pathogen surveillance system. Nature Reviews Genetics, 19(1), 9–20. https://doi.org/10.1038/nrg.2017.88

Grossman, R. L. (2019). Data Lakes, Clouds, and Commons: A Review of Platforms for Analyzing and Sharing Genomic Data. Trends in Genetics: TIG, 35(3), 223–234. https://doi.org/10.1016/j.tig.2018.12.006

Grzesik, P., Augustyn, D. R., Wyciślik, Ł., & Mrozek, D. (2021). Serverless computing in omics data analysis and integration. Briefings in Bioinformatics, bab349. https://doi.org/10.1093/bib/bbab349

Hawari, M. A., Hong, C. S., & Biesecker, L. G. (2021). SomatoSim: Precision simulation of somatic single nucleotide variants. BMC Bioinformatics, 22, 109. https://doi.org/10.1186/s12859-021-04024-8

Koppad, S., B, A., Gkoutos, G. V., & Acharjee, A. (2021). Cloud Computing Enabled Big Multi-Omics Data Analytics. Bioinformatics and Biology Insights, 15, 11779322211035920. https://doi.org/10.1177/11779322211035921

Krissaane, I., De Niz, C., Gutiérrez-Sacristán, A., Korodi, G., Ede, N., Kumar, R., Lyons, J., Manrai, A., Patel, C., Kohane, I., & Avillach, P. (2020). Scalability and cost-effectiveness analysis of whole genome-wide association studies on Google Cloud Platform and Amazon Web Services. Journal of the American Medical Informatics Association: JAMIA, 27(9), 1425–1430. https://doi.org/10.1093/jamia/ocaa068

Krusche, P., Trigg, L., Boutros, P. C., Mason, C. E., De La Vega, F. M., Moore, B. L., Gonzalez-Porta, M., Eberle, M. A., Tezak, Z., Lababidi, S., Truty, R., Asimenos, G., Funke, B., Fleharty, M., Chapman, B. A., Salit, M., Zook, J. M., & Global Alliance for Genomics and Health Benchmarking Team. (2019). Best practices for benchmarking germline small-variant calls in human genomes. Nature Biotechnology, 37(5), 555–560. https://doi.org/10.1038/s41587-019-0054-x

Langmead, B., & Nellore, A. (2018). Cloud computing as a platform for genomic data analysis and collaboration. Nature Reviews. Genetics, 19(4), 208–219. https://doi.org/10.1038/nrg.2017.113

Larson, D. E., Harris, C. C., Chen, K., Koboldt, D. C., Abbott, T. E., Dooling, D. J., Ley, T. J., Mardis, E. R., Wilson, R. K., & Ding, L. (2012). SomaticSniper: Identification of somatic point mutations in whole genome sequencing data. Bioinformatics, 28(3), 311–317. https://doi.org/10.1093/bioinformatics/btr665

Leonard, C., Wood, S., Holmes, O., Waddell, N., Gorse, D., Hansen, D. P., & Pearson, J. V. (2019). Running Genomic Analyses in the Cloud. Studies in Health Technology and Informatics, 266, 149–155. https://doi.org/10.3233/SHTI190787

Li, H., Handsaker, B., Wysoker, A., Fennell, T., Ruan, J., Homer, N., Marth, G., Abecasis, G., Durbin, R., & 1000 Genome Project Data Processing Subgroup. (2009). The Sequence Alignment/Map format and SAMtools. Bioinformatics, 25(16), 2078–2079. https://doi.org/10.1093/bioinformatics/btp352

Liu, B., Madduri, R. K., Sotomayor, B., Chard, K., Lacinski, L., Dave, U. J., Li, J., Liu, C., & Foster, I. T. (2014). Cloud-based bioinformatics workflow platform for large-scale next-generation sequencing analyses. Journal of Biomedical Informatics, 49, 119–133. https://doi.org/10.1016/j.jbi.2014.01.005

Mölder, F., Jablonski, K. P., Letcher, B., Hall, M. B., Tomkins-Tinch, C. H., Sochat, V., Forster, J., Lee, S., Twardziok, S. O., Kanitz, A., Wilm, A., Holtgrewe, M., Rahmann, S., Nahnsen, S., & Köster, J. (2021). Sustainable data analysis with Snakemake. F1000Research, 10, 33. https://doi.org/10.12688/f1000research.29032.2

Nwadiugwu, M. C., & Monteiro, N. (2022). Applied genomics for identification of virulent biothreats and for disease outbreak surveillance. Postgraduate Medical Journal. https://doi.org/10.1136/postgradmedj-2021-139916

Poplin, R., Chang, P.-C., Alexander, D., Schwartz, S., Colthurst, T., Ku, A., Newburger, D., Dijamco, J., Nguyen, N., Afshar, P. T., Gross, S. S., Dorfman, L., McLean, C. Y., & DePristo, M. A. (2018). A universal SNP and small-indel variant caller using deep neural networks. Nature Biotechnology, 36(10), 983–987. https://doi.org/10.1038/nbt.4235

Poplin, R., Ruano-Rubio, V., DePristo, M. A., Fennell, T. J., Carneiro, M. O., Auwera, G. A. V. der, Kling, D. E., Gauthier, L. D., Levy-Moonshine, A., Roazen, D., Shakir, K., Thibault, J., Chandran, S., Whelan, C., Lek, M., Gabriel, S., Daly, M. J., Neale, B., MacArthur, D. G., & Banks, E. (2018). Scaling accurate genetic variant discovery to tens of thousands of samples (p. 201178). bioRxiv. https://doi.org/10.1101/201178

Ray, U., Krishnan, V., Bahmani, A., Pan, C., Bettinger, K., Tsao, P., Mueller, F., & Snyder, M. (2021). Hummingbird: Efficient Performance Prediction for Executing Genomics Applications in the Cloud. Bioinformatics, 37(17), 2537–2543.

Rosati, S. (2020). Comparison of CPU and Parabricks GPU Enabled Bioinformatics Software for High Throughput Clinical Genomic Applications. Master’s Thesis (2009-), 43.

Tanjo, T., Kawai, Y., Tokunaga, K., Ogasawara, O., & Nagasaki, M. (2021). Practical guide for managing large-scale human genome data in research. Journal of Human Genetics, 66(1), 39–52. https://doi.org/10.1038/s10038-020-00862-1

Van der Auwera, G. A., & O’Connor, B. D. (2020). Genomics in the cloud: Using Docker, GATK, and WDL in Terra (1st ed.). O’Reilly Media.

Zhang, Q., Liu, H., & Bu, F. (2021). High performance of a GPU-accelerated variant calling tool in genome data analysis [Preprint]. Bioinformatics. https://doi.org/10.1101/2021.12.12.472266

Zhao, S., Agafonov, O., Azab, A., Stokowy, T., & Hovig, E. (2020). Accuracy and efficiency of germline variant calling pipelines for human genome data (p. 2020.03.27.011767). bioRxiv. https://doi.org/10.1101/2020.03.27.011767

Zook, J. M., Catoe, D., McDaniel, J., Vang, L., Spies, N., Sidow, A., Weng, Z., Liu, Y., Mason, C. E., Alexander, N., Henaff, E., McIntyre, A. B. R., Chandramohan, D., Chen, F., Jaeger, E., Moshrefi, A., Pham, K., Stedman, W., Liang, T., … Salit, M. (2016). Extensive sequencing of seven human genomes to characterize benchmark reference materials. Scientific Data, 3(1), 160025. https://doi.org/10.1038/sdata.2016.25

