## Supplemental Information for "Accelerating genomic workflows using NVIDIA Parabricks"

**Table S1.** Results of benchmarking for in AWS with p4 machine family with A100 GPUs.

| Platform | Pipeline | VM-Type | Variant-Caller | Time (min) | Time (hours) | Cost ($) | Fold Acceleration | % Cost-Savings |
| --- | --- | --- | --- | --- | --- | --- | --- | --- |
| AWS | Germline | C6i.8xlarge | DeepVariant | 1317.3 | 21.96 | 29.9 | _ | _ |
|  |  | GPU2 |  | 116.7 | 1.94 | 63.7 | 11.3 | -113.4 |
|  |  | GPU4 |  | 61.2 | 1.02 | 33.4 | 21.5 | -11.9 |
|  |  | GPU8 |  | 42.9 | 0.72 | 23.5 | 30.7 | 21.5 |
| AWS | Germline | C6i.8xlarge | HaplotypeCaller | 2175.9 | 36.26 | 49.32 | _ | _ |
|  |  | GPU2 |  | 85.6 | 1.4 | 46.8 | 25.4 | 5.2 |
|  |  | GPU4 |  | 49.2 | 0.82 | 26.9 | 44.2 | 45.5 |
|  |  | GPU8 |  | 33.4 | 0.56 | 18.3 | 65.1 | 63 |
| AWS | Somatic | C6i.8xlarge | LoFreq | 180.2 | 3 | 4.1 | _ | _ |
|  |  | GPU2 |  | 114.9 | 1.92 | 62.8 | 1.6 | -1437.2 |
|  |  | GPU4 |  | 72.2 | 1.20 | 39.4 | 2.5 | -865.3 |
|  |  | GPU8 |  | 49.1 | 0.82 | 26.8 | 3.7 | -557.1 |
|  |  | GPU8 |  | 49.5 | 0.83 | _ | _ | _ |
| AWS | Somatic | C6i.8xlarge | Muse | 425.1 | 7.09 | 9.6 | _ | _ |
|  |  | GPU2 |  | 29.7 | 0.49 | 16.2 | 14.3 | -68.1 |
|  |  | GPU4 |  | 22.5 | 0.37 | 12.3 | 18.9 | -27.5 |
|  |  | GPU8 |  | 27.5 | 0.46 | 15 | 15.5 | -55.8 |
| AWS | Somatic | C6i.8xlarge | Mutect2 | 414.51 | 6.91 | 9.40 | _ | _ |
|  |  | GPU2 |  | 18.8 | 0.31 | 10.3 | 22.06 | -9.25 |
|  |  | GPU4 |  | 17.89 | 0.30 | 9.8 | 23.17 | -4 |
|  |  | GPU8 |  | 28 | 0.47 | 15.3 | 14.80 | -62.78 |
| AWS | Somatic | C6i.8xlarge | SomaticSniper | 391.9 | 6.53 | 8.88 | _ | _ |
|  |  | GPU2 |  | 52.8 | 0.88 | 28.9 | 7.4 | -224.9 |
|  |  | GPU4 |  | 50.8 | 0.85 | 27.7 | 7.7 | -212.4 |
|  |  | GPU8 |  | 93.4 | 1.56 | 51 | 0.6 | -474.6 |

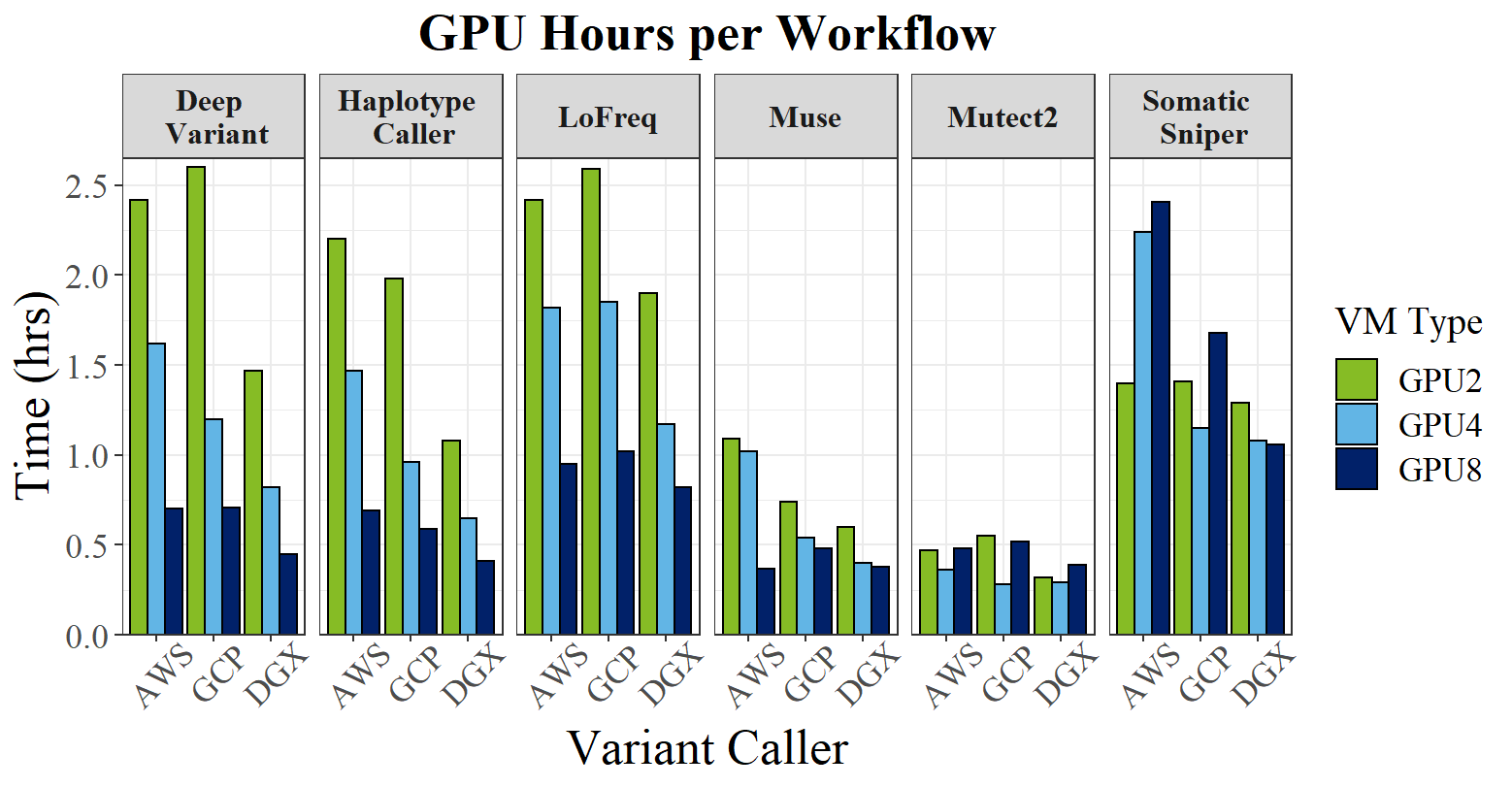

***Figure S1****: GPU benchmarking results for AWS, GCP, and NVIDIA DGX with the V100 GPUs on AWS.*

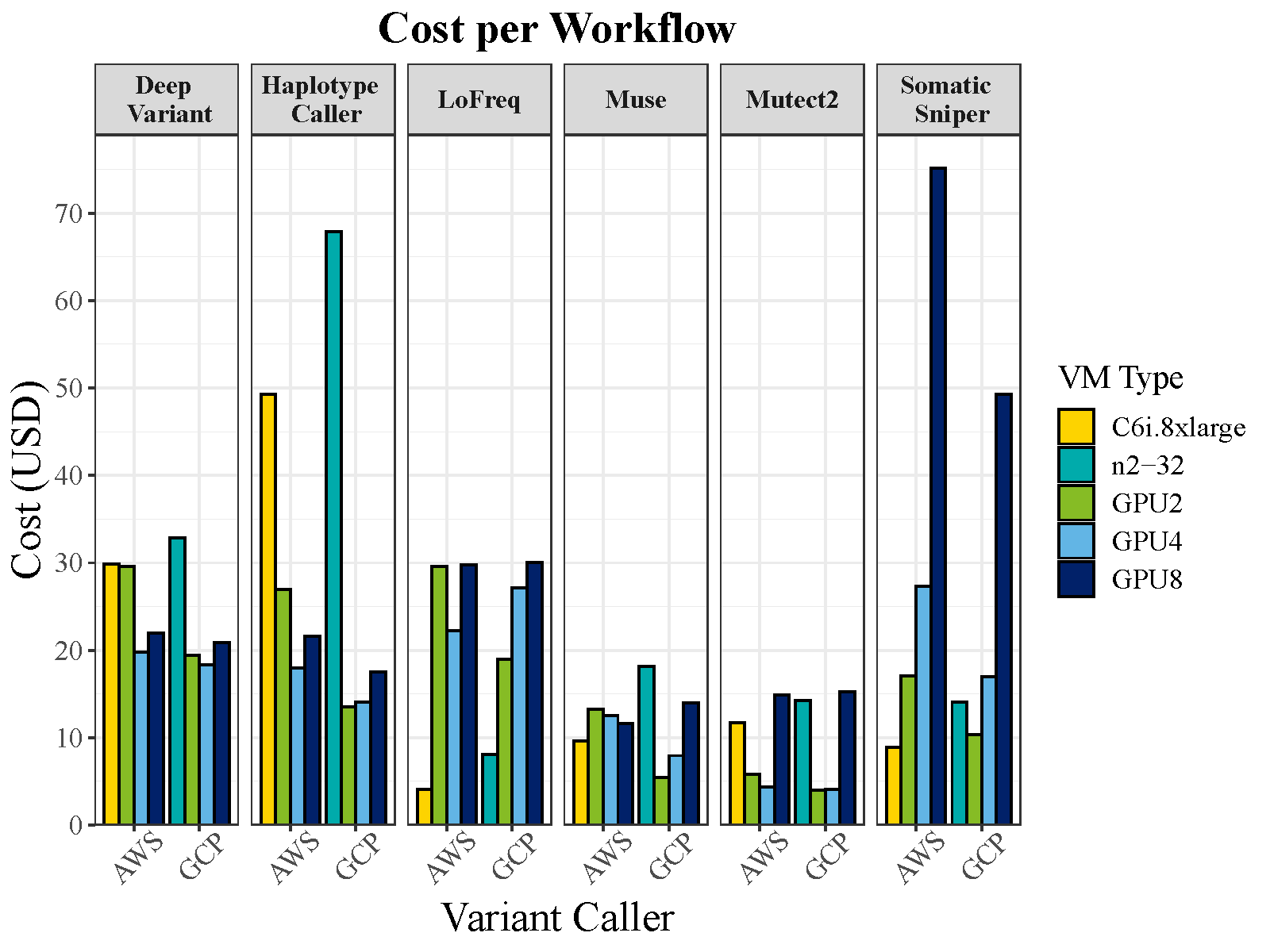

***Figure S2****: Cost per workflow (V100 machine on AWS) comparing CPU to GPU costs for each machine type.*

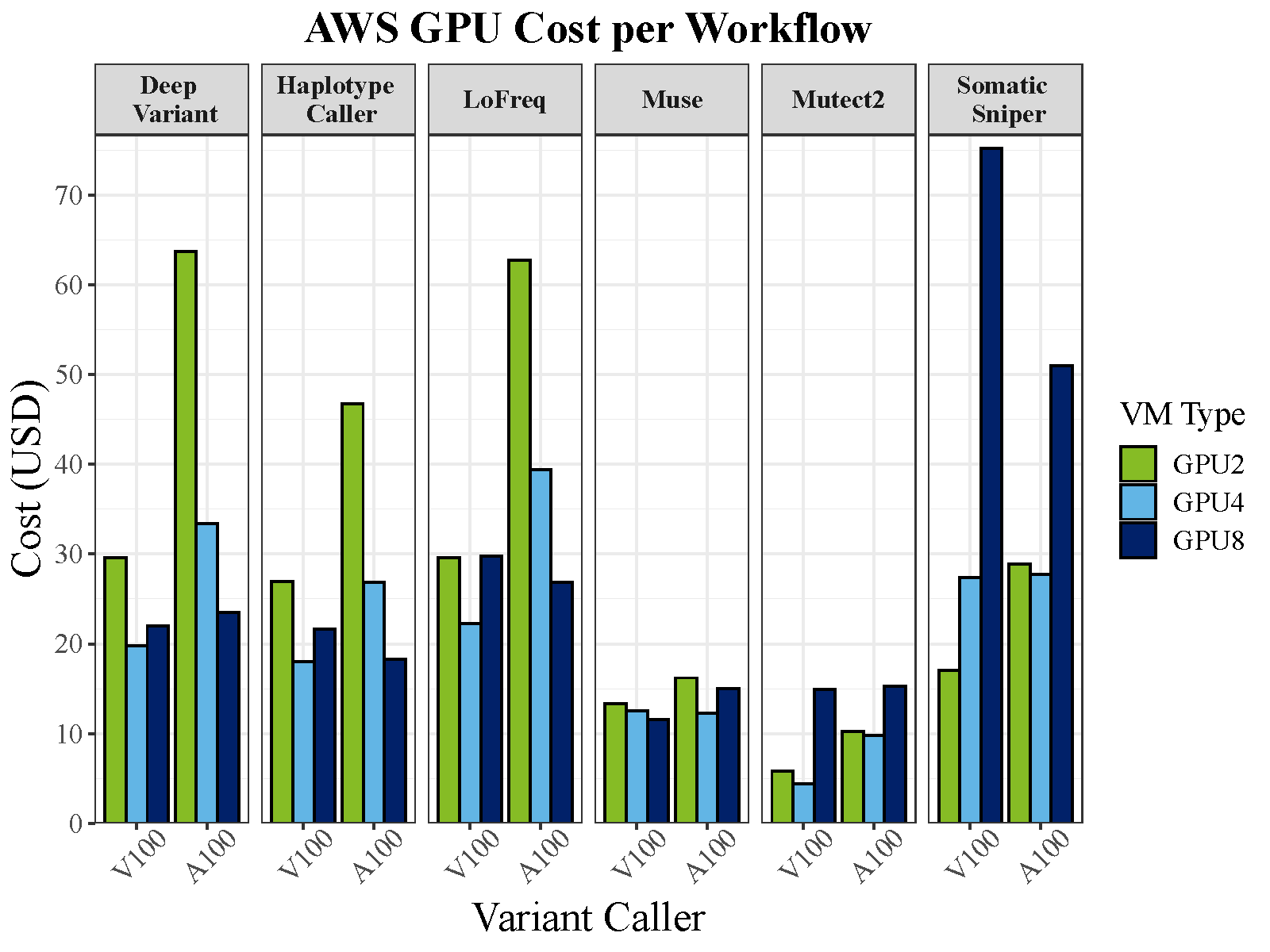

***Figure S3****: Cost differences between A100 and V100 GPU machines on AWS.*

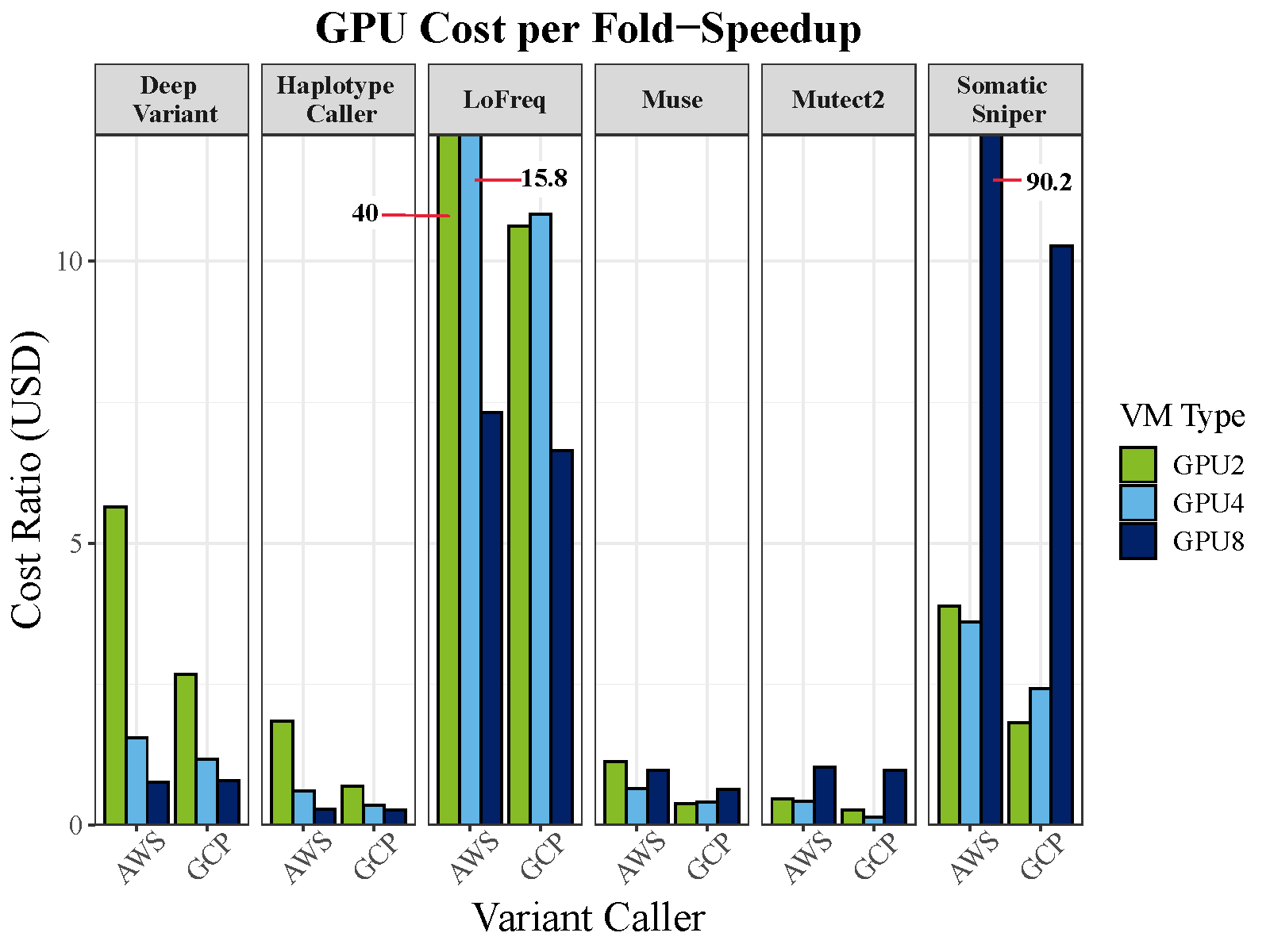

***Figure S4.*** *Comparison of AWS (A100 machine) vs. GCP GPU cost ratio per variant caller. Cost ratio being the ratio between cost per hour and fold speed-up. Cost per fold-speedup shows the benefit of harnessing GPU over CPU in select algorithms, while other algorithms are more cost-efficient with CPUs.*

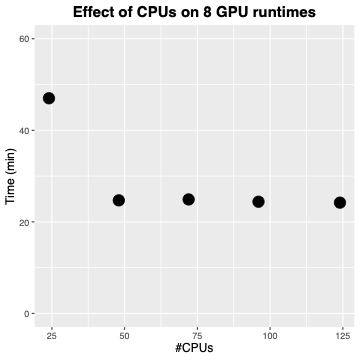

Figure S5: HaplotypeCaller runtime (in minutes) with 8 A100 GPUs on the NVIDIA DGX plotted by number of CPUs allocated per run. X axis shows runs with 24, 48, 74, 96, and 124 CPUs.
